# Stable behavioral and neural responses to thermal stimulation despite large changes in the *Hydra vulgaris* nervous system

**DOI:** 10.1101/787648

**Authors:** Constantine N. Tzouanas, Soonyoung Kim, Krishna N. Badhiwala, Benjamin W. Avants, Jacob T. Robinson

## Abstract

Many animals that lose neural tissue due to injury or disease have the ability to maintain their behavioral abilities by regenerating new neurons or reorganizing existing neural circuits. However, most small model organisms used for neuroscience like nematodes and flies lack this high degree of neural plasticity. These animals often show significant behavioral deficits if they lose even a single neuron. Here we show that the small freshwater cnidarian *Hydra vulgaris* can maintain stable sensory motor behaviors even after losing half of the neurons in its body. Specifically, we find that both the behavioral and neural response to a rapid change in temperature is maintained if we make their nervous system roughly 50% smaller by caloric restriction or surgery. These observations suggest that *Hydra* provides a rich model for studying how animals maintain stable sensory-motor responses within dynamic neural circuit architectures, and may lead to general principles for neural circuit plasticity and stability.

**Significance Statement:** The ability of the nervous system to restore its function following injury is key to survival for many animals. Understanding this neural plasticity in animals across the phylogenetic tree would help reveal fundamental principles of this important ability. To our knowledge, the discovery of a set of neurons in the jellyfish polyp *Hydra vulgaris* that stably support a response to thermal stimulation is the first demonstration of large-scale neural plasticity in a genetically tractable invertebrate model organism. The small size and transparency of *Hydra* suggests that it will be possible to study large-scale neural circuit plasticity in an animal where one can simultaneously image the activity of every neuron.

## Introduction

One remarkable feature of the nervous system is plasticity - the ability to alter or reorganize neural circuits to gain or restore function. In mammals, individual neurons can alter their excitability based on presynaptic transmission frequencies [1] to maintain stable activity levels. Deficits in this process of homeostatic plasticity have been associated with neural disorders [2]. Similarly, synaptic plasticity allows collections of neurons to remap connections to produce behavioral compensation after broad injuries such as strokes, using molecular mechanisms similar to those involved in neural development [3]. Despite these insights, general principles for how neural plasticity helps to preserve behavioral function are not well known across phylogeny.

Small invertebrates would be ideal model systems for studying the processes associated with neural plasticity and behavior because they have relatively small nervous systems that make it easier to track neural circuit reorganization in living animals. However, most invertebrates show modest regeneration or lack well-developed transgenic techniques. For instance, even weeks after nerve lesions, *Aplysia* shows incomplete recovery of the number of large axons in nerve cores [4], receptive fields [5], and behaviors like head waving during feeding [6] and tail-elicited siphon-withdrawal reflex [7]. Neurons in the central nervous system of *Drosophila* do not regenerate following axotomies in either whole-brain cultures [8] or in larvae [9]. In *C. elegans*, axons of select neurons can regenerate after being severed [10]; however, growth trajectories are error-prone [11] even when identical neurons across animals are subjected to similar injury protocols [12]. Exceptional neural plasticity can be found in planarians, which can reform entire organisms [13]; from small fragments of tissue, but these animals lack a suite of transgenic tools similar to that of *Drosophila* and *C. elegans* [14].

*Hydra vulgaris* is unique among invertebrate model organisms in that they are amenable to a variety of transgenic techniques and display significant neural plasticity. As examples of *Hydra*’s remarkable regenerative capabilities, it can regrow its entire body and nervous system from a fragment as small as ~300 cells [15], [16], and its entire nervous system can be rebuilt from even a single stem cell [17]. The regenerative properties of *Hydra*’s nervous system are due to the multipotent interstitial stem cells which continually give rise to several cell types, including the neurons. Even in an uninjured animal, there is continual turnover of differentiated neurons, with all neurons being replaced every 20 days [14]. These properties of the *Hydra* nervous system make it possible to study how new neurons integrate into existing circuits and how neuronal circuits are rebuilt following injury.

The small size and genetic tractability of *Hydra* provide additional advantages. As a millimeter-scale invertebrate, an entire *Hydra* can be imaged at single-cell resolution [15]. Furthermore, the ability to generate transgenic lines by embryo injection [18], [19] has enabled functional calcium imaging in neurons [20] and epitheliomuscular cells [21]. In addition, cell-type specific promoters have been developed based on single-cell transcriptomic analysis of *Hydra* that revealed twelve neuronal subtypes, each expressing distinct sets of biomarkers linked to neuronal development and function [22], [23].

Given *Hydra’s* potential as a model for neural plasticity, it is critical to establish behavioral assays that indicate if and when neural circuits have regained their ability to regulate behavior. While *Hydra* has been studied for over 300 years, we lack quantitative characterization of behaviors like sensory-motor responses. Studies of behavior dating back to the 1700’s [24] qualitatively describe responses to light [25], chemicals [26], and temperature [27]–[29], but these experiments fall short of quantitative evaluation of behavior and neural activity. Recently, machine learning approaches have been used to identify behavioral motifs in freely moving *Hydra*, but these experiments have not been extended to sensory-motor responses [30].

*Hydra’s* response to a rapid change in temperature is one sensory-motor behavior that could be used to assess neural plasticity, but it must first be precisely defined and quantified. While prior experiments suggest that *Hydra* can sense temperature, more work is needed to quantitatively characterize the animal’s neural and behavioral response patterns to a rapid temperature change. Mast showed that when touched by heated objects, *Hydra* responds by bending toward the stimulus, but these experiments cannot fully distinguish between mechanosensory and thermosensory behaviors [28]. Schroeder and Callaghan reported the highest and lowest temperatures in which *Hydra oligactis* and *Hydra pseudoligactis* could survive [27]. Bosch et al. demonstrated that exposing *Hydra* to moderately elevated temperatures (30°C for two hours) increases their ability to survive culture at high temperatures (34°C for four days) that would otherwise be lethal [29]. These experiments show that *Hydra* viability depends on the ambient temperature, but they do not reveal the specific sensory-motor response to an acute thermal stimulus.

Here we show the first quantitative description of *Hydra’s* behavioral and neural response to thermal stimulation and demonstrate that these responses are maintained even if the numbers of neurons in the animal change by a factor two due to caloric restriction or surgery. To perform this study, we developed a microfluidic device capable of delivering rapid and precise thermal stimuli without the confound of mechanical stimulation. Using this technique, we find that *Hydra* elongates after the onset of positive thermal stimulation, followed by a contraction lasting for the stimulus duration. Synchronous with body movements, we find temperature-dependent oscillatory neural activity within a ring of neurons in *Hydra*’s peduncle. The frequency of this oscillation decreases for negative thermal stimulation and increases for positive thermal stimulation. We show that the frequency of the neural oscillation depends primarily on the absolute temperature of the thermal stimuli, rather than relative changes from culture baseline. Importantly, we find that these frequencies are nearly unchanged if the number of neurons in these animals change by a factor of two, whether due to caloric restrictions or surgical sectioning. These results suggest that the *Hydra*’s nervous system is a valuable model for studying neural plasticity and robust neural circuit architectures.

## Results

### Microfluidic device enables thermal stimulation and whole-body imaging of *Hydra vulgaris*

Because *Hydra* are sensitive to mechanical stimulation (e.g., touch and changes in fluid flow [31],[32]), we designed a two-layer microfluidic device to deliver thermal stimuli without mechanical confounds: a *Hydra* immobilization chamber fabricated above a flow layer with multiple inlet ports (Fig. 1a, see Materials and Methods). By connecting the flow layer’s inlet ports to in-line heaters at different temperature setpoints, we can provide rapid and precise thermal stimulation by switching fluid flow through the different inlet ports – a process that we automate using microcontrollers (see Materials and Methods). In our experiments, one inlet (i in Fig. 1a) provides fluid that is maintained at *Hydra*’s culture temperature to serve as a control. A second inlet provides fluid that is maintained at a stimulus temperature ranging from 9°C to 36°C. To validate our ability to rapidly and repeatedly modulate the temperature of the *Hydra* immobilization chamber, we measured the temperature of the device glass with an IR camera (Fig. 1b-c). We found that we could switch between temperature setpoints in approximately one second, and that the temperature set points were repeatable to within a standard deviation of 0.5°C across multiple stimulation cycles.

**Figure 1:**
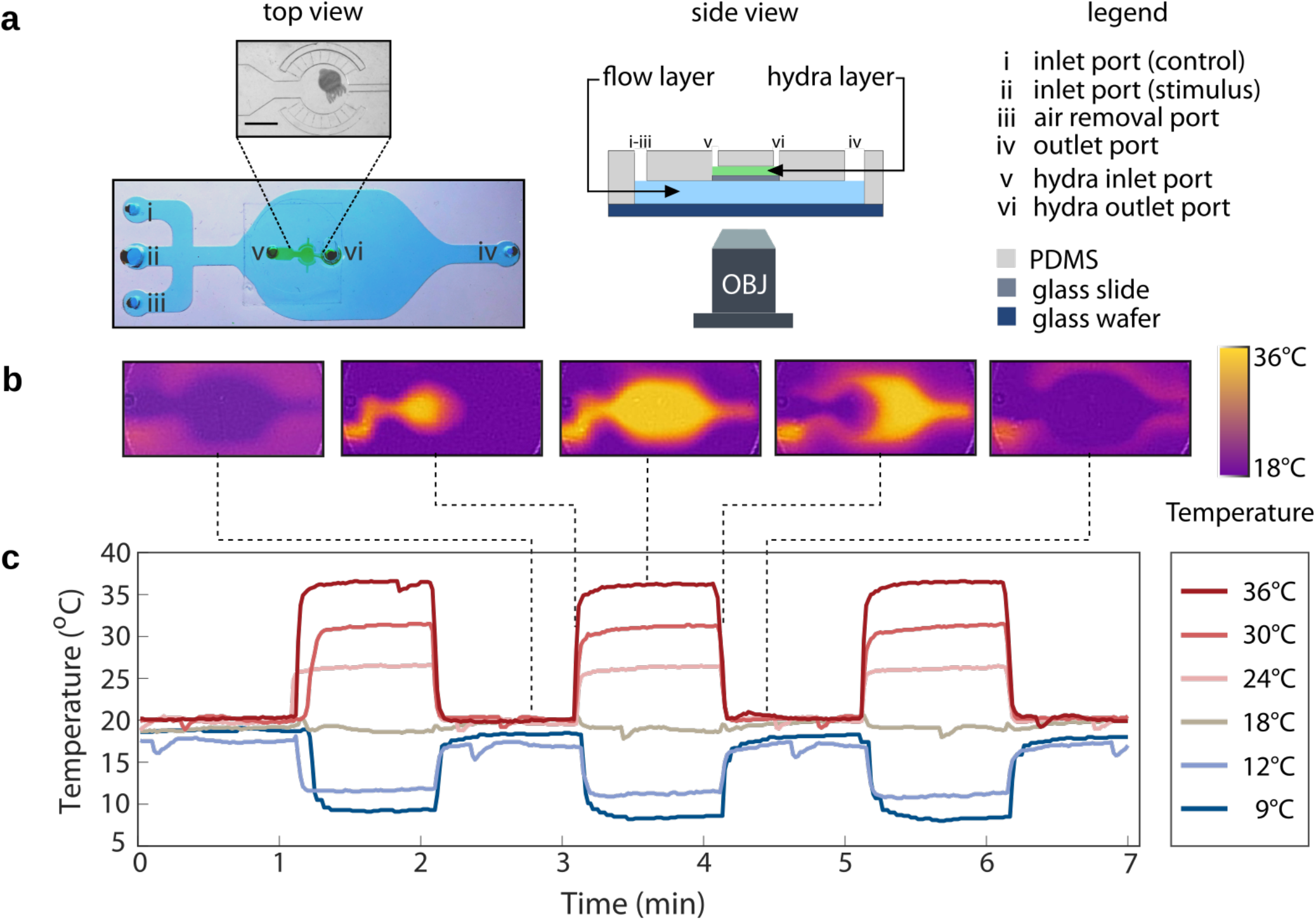
Characterization and validation of two-layer microfluidic device for thermal stimulation of *Hydra vulgaris*. **a** (left) Photograph of microfluidic device with blue dye in flow layer and green dye in *Hydra* chamber. Scale bar = 1 mm. (middle) Schematic of cross-section of the device. (right) legend for left and middle panel. **b** Infrared image showing temperature of the flow layer before (leftmost), the start of (second leftmost), during (center), end of (second rightmost), and after (rightmost) thermal stimulation. **c** Time course of thermal stimulation at temperatures above and below *Hydra*’s baseline culture temperature of 18°C.

### Positive thermal stimulation drives sequential elongation and contraction in *Hydra vulgaris*

Having fabricated a microfluidic device that allows us to thermally stimulate and simultaneously image *Hydra*, we supplied fluids of different temperatures to the flow layer to measure how *Hydra* responds to a rapid change in temperature. We observed no statistically significant changes to *Hydra*’s body length in experiments when we switched the source of the flow layer between two sources at 18°C compared to experiments with no fluid flow (“18°C” and “No flow” curves in Fig. 2C), showing the developed microfluidic device does not produce mechanical stimulation.

**Figure 2:**
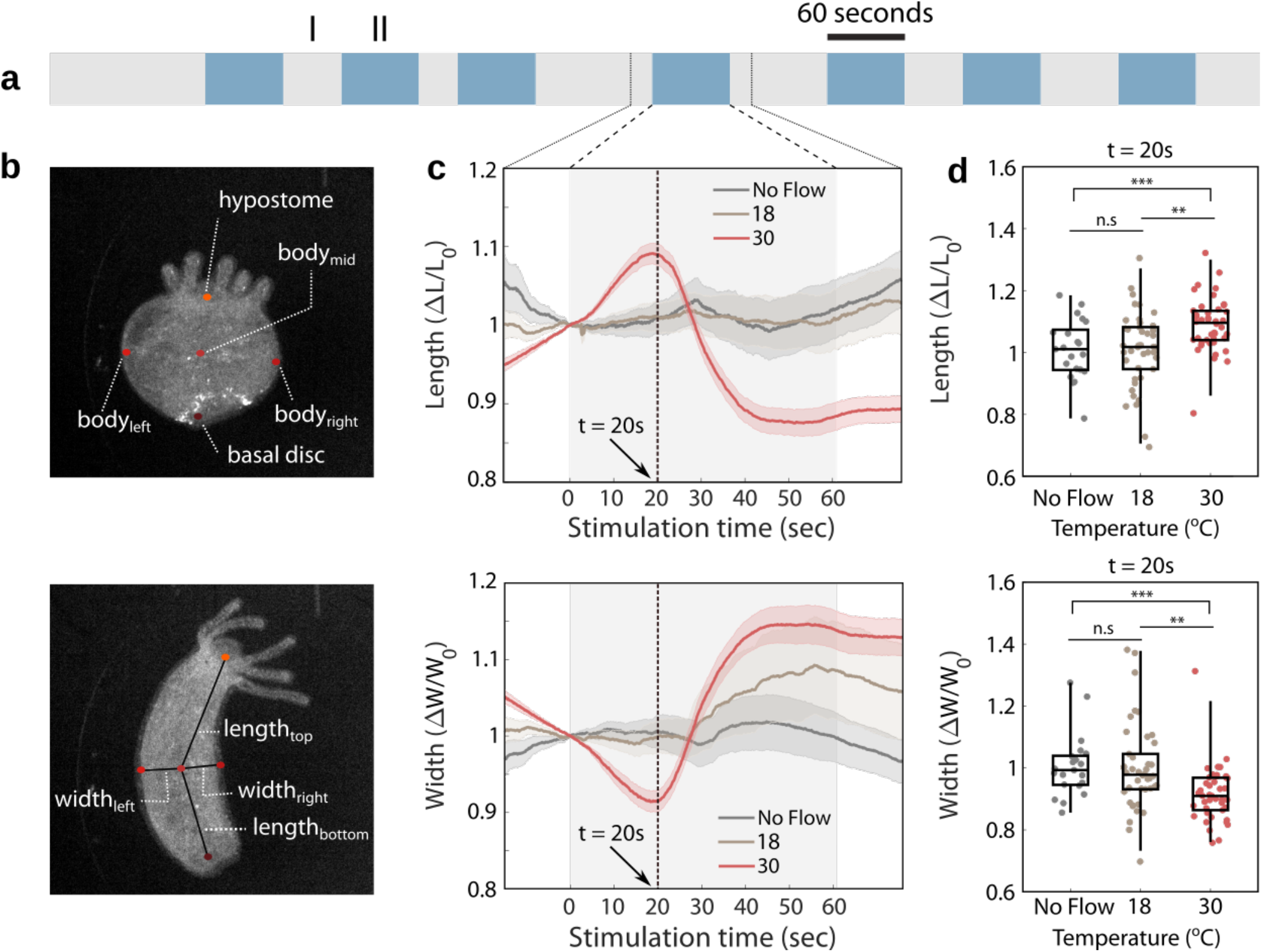
*Hydra*’s body length initially increases and width initially decreases when stimulated at 30°C. **a** Schematic of a representative thermal stimulation protocol. Gray regions (I) indicate non-stimulus periods at *Hydra*’s culture temperature lasting between 30 and 90 seconds, and blue-colored regions (II) indicate stimulus periods at designated temperatures for 60 seconds. **b** Representative labeled frame from DeepLabCut, annotated. Top panel describes which body points were annotated, and the bottom panel shows how the length and width of the animals are determined. The hypostome is the end of the body with *Hydra*’s mouth and tentacles (oral end), while the basal disk is used to adhere to substrates (aboral end). **c** Average length and width of *Hydra* calculated for stimulation periods and time-aligned to stimuli, with 15 seconds before and after stimulation periods. Shaded error bars correspond to standard error. **d** The change in length (top panel) and width (bottom panel) at 20 seconds after onset of stimulation, for each stimulus period (t = 20s). N=3 *Hydra* for No Flow, N=6 *Hydra* for 18°C and 30°C. (n.s = not significant, ** p<0.001, *** p<0.0001, unpaired t-test).

To avoid synchronizing thermal stimuli with natural rhythmic behavior [32], [33] or entraining a periodic response, we applied the thermal stimuli for a period of 60 seconds and randomized the time interval between successive stimuli (Fig. 2a). To quantify changes in *Hydra*’s posture, we applied DeepLabCut [34] to automate labeling of *Hydra*’s body width and length using hypostome (oral end), basal disc (aboral end), leftmost, middle, and rightmost points in the body column region (Fig. 2b, see Materials and Methods). We found *Hydra* respond to a rapid change in temperature (18°C to 30°C over approximately 1s) by first elongating, and then longitudinally contracting. The peak body length occurs approximately 20 seconds after the stimulus onset (Fig. 2c-d).

### Subset of peduncle neurons show periodic calcium spikes in response to thermal stimulation

Using fluorescent imaging of transgenic *Hydra* that express the calcium-sensitive fluorophore GCaMP6s in neurons, we observed bursts of synchronous calcium spikes following thermal stimulation in a group of neurons in the animal’s peduncle (Fig. 3b). We will refer to these neurons as temperature responsive (TR) neurons. To quantify this effect, we selected a region of interest that encompassed the entire *Hydra* peduncle (Fig. 3b). Although not all neurons in this region are putative TR neurons, the strong signal produced by the synchronous TR neuron activity allowed us to measure the calcium spike rate using this region of interest (ROI) (Fig. 3c-d, Supplemental Video 1-2).

**Figure 3:**
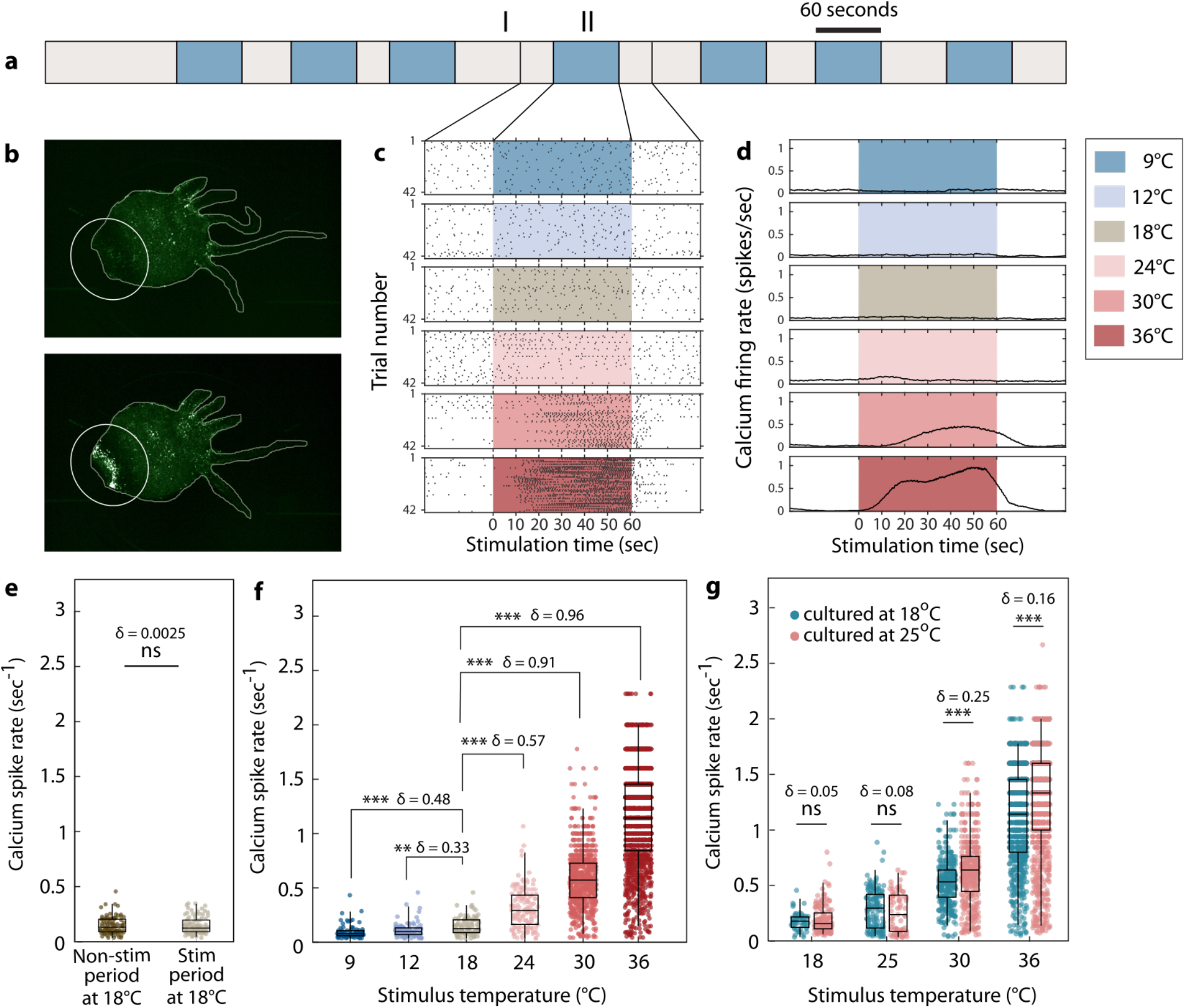
Thermal stimulation modulates the synchronous neural activity in the peduncle of *Hydra vulgaris*. **a** Schematic of a representative thermal stimulation protocol. Gray-colored regions (I) indicate non-stimulus periods at *Hydra*’s culture temperature (either 18°C or 25°C) lasting between 30 and 90 seconds. The blue-colored regions (II) indicate stimulus periods at designated temperatures for 60 seconds. **b** Images of *Hydra* without synchronous neural firing (top) and during firing (bottom) in the peduncle (white circle). **c** Raster plot of synchronous firing events in peduncle in **b**. One black vertical line indicates one synchronous firing event. Each row represents the raster plot of spikes time-aligned with one stimulation (boxed regions) with 30 seconds before and after stimulation (white non-boxed region). **d** Peristimulus time histogram (calcium firing rate) of **c** which was calculated with a 10 second sliding window. (ns = not significant, Mann-Whitney U Test. δ = Effect size using Cliff’s delta) **e** Calcium spike rate comparison between the non-stimulation (gray-colored region I in **a**) and stimulation (blue-colored region II in **a**) periods with a stimulus temperature of 18°C. **f** Calcium spike rates from large and small *Hydra*, all cultured at 18°C. N=3 large animals; N=3 small animals. (ns = not significant, ** p<0.0001, *** p<0.00001, Mann-Whitney U Test. δ = Effect size using Cliff’s delta). **g** Calcium spike rates from Hydra cultured at 18°C and 25°C. X-axis notes the stimulation temperatures. N=3 for all animals. (ns = not significant, *** p<0.00001, Mann-Whitney U test. δ = Effect size using Cliff’s delta)

We found that the calcium spike rate did not significantly change when switching between fluid reservoirs at 18°C, suggesting that changing fluids in the flow layer did not produce a mechanical stimulation artifact (Fig. 3e, effect size δ = 0.0025). When we thermally stimulated *Hydra* across a wide range of temperatures both above and below their culture temperature, we observed that the TR neurons’ calcium spike rate during positive thermal stimulation (i.e., heating) was significantly higher than in control experiments (compare conditions labeled “24°C”, “30°C”, and “36°C” to the control condition of “18°C” in Fig. 3c-d,f). Likewise, TR neurons’ firing rate decreased during negative thermal stimulation (i.e., cooling; compare conditions labeled “9°C” and “12°C” to the control condition of “18°C” in Fig. 3c-e).

We also found that the firing rate of the TR neurons during thermal stimulation depends primarily on the absolute temperature of the thermal stimulus and not on the relative increase in temperature. We found that when we increased the *Hydra* culture temperature from 18°C to 25°C, there was no significant change in the calcium spike rate for most of the thermal stimulus temperatures. At high stimulus temperatures (i.e., 30°C and 36°C in Fig. 3g) we observed a statistically significant difference based on culture temperature, but the effect size was small (δ = 0.25 at 30°C stimulation, δ = 0.16 at 36°C stimulation) compared to changing the stimulation temperature by a comparable amount (e.g., δ = 0.71 between *Hydra* stimulated at 30°C or 36°C, but both cultured at 18°C). These results support the conclusion that the calcium spike rate of TR neurons depends primarily on the absolute temperature of a thermal stimulus.

### *Hydra vulgaris*’ neural response to thermal stimulation is robust against naturally-occurring two-fold changes in neuron count

Having identified that the calcium spike rate of TR neurons encodes the absolute temperature of a thermal stimulus, we asked if this encoding is maintained even if animals have significantly different numbers of neurons. To answer this question, we first determined that the number of neurons in *Hydra* is linearly related to the size of the animal. Using longitudinal fluorescence imaging of *Hydra* over two weeks of starvation or regular feedings, we found that the number of interstitial cells (known to be comprised of 58.9% neurons, 23.8% progenitor cells, 7.8% nematocytes/nematoblasts, and 0.4% germline cells [22]) scaled with *Hydra*’s body volume, consistent with prior reports of the density of neurons as a function of animal size [35], [36] (Fig. 4a-c).

**Figure 4:**
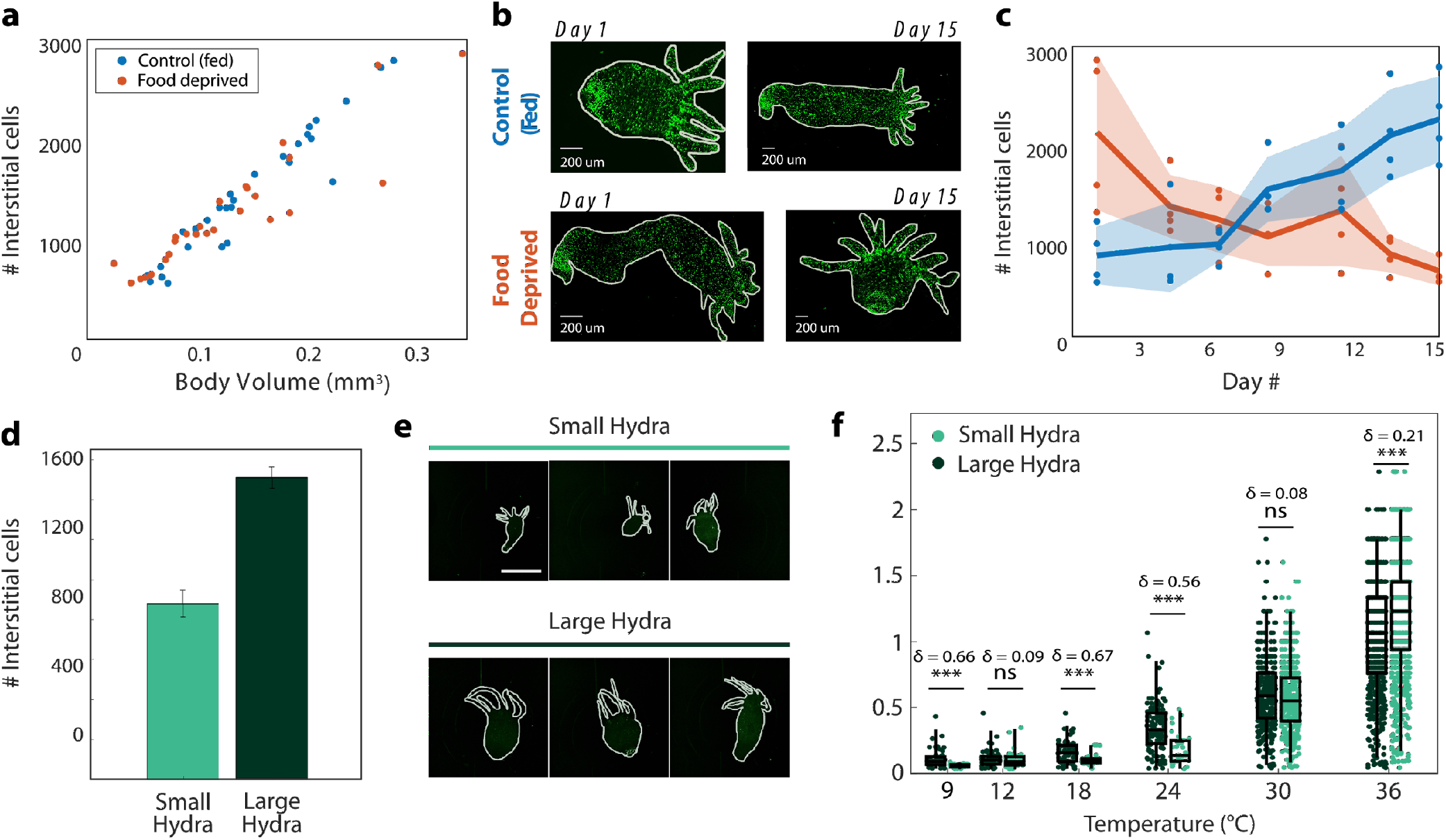
Frequency of thermal response neurons does not depend on the number of neurons in *Hydra*. **a**Number of interstitial cells (comprising 58.9% neurons, 23.8% progenitor cells, 7.8% nematocytes/nematoblasts, and 0.4% germline cells) as a function of animal size. **b** Fluorescent image of Hydra at days 1 and 15 for both control and food deprived groups (pseudo colored). **c** Change in the number of interstitial cells over the time course of 15 days. **d** Estimated number of interstitial cells from measurements in **a**. **e** Three representative *Hydra* from small and large groups (pseudo colored) Scale bar = 500 μm. **f** Spike rates from large (N=3) and small (N=3) *Hydra*, all cultured at 18°C. (ns = not significant, *** p<0.00001, Mann-Whitney U test. δ = Effect size using Cliff’s delta)

To compare the thermal responses of animals with different sizes, we separated animals into “large” and “small” cohorts according to their body size. Based on the differences in body size and our measured relationship between body size and number of neurons, we estimate that the difference in neuron count between these groups is approximately two-fold (Fig. 4d-e). Note that while the small and large populations have a two-fold difference in the mean, the distributions do overlap due to the naturally occurring inhomogeneity in the small and large populations (Supplemental Fig. 1). Nevertheless, we expect that these groups would show differences in behavior if the thermal response were dependent upon the number of neurons in the animal.

When we measured the calcium spike rates during thermal stimulation, we found a nearly identical temperature dependence in both small and large animals (Fig. 4f), suggesting that *Hydra* maintain stable encoding of thermal stimuli despite significant, two-fold differences in the size of their neural circuits. Both groups exhibit decreased calcium spike rates during negative thermal stimulation and increased calcium spike rates during positive thermal stimulation. Although some temperatures did show statistically significant differences between the thermal response of the “small”; and “large” animals (Fig. 4f), the effect size (δ) is small compared to the effect size of changes in stimulus temperature (e.g., δ = 0.96 when stimulating at 36°C as compared to culture temperature baseline; δ = 0.21 between large and small animals when stimulated at 36°C). At 9°C stimulation and control temperature (18°C), the effect size δ between the small and large group was 0.66 and 0.67, respectively, which is comparable to the effect size of when comparing the calcium spike rate from the same animal measured on subsequent days (δ = 0.61, Supplemental Figure 1). Despite remodeling their nervous system to account for a two-fold increase or decrease in the numbers of neurons, *Hydra* are able to consistently encode the temperature of a thermal stimulus.

### Surgical removal of half of *Hydra*’s neurons does not significantly affect neural response to thermal stimulation

While animals with naturally-occurring large and small nervous systems showed similar responses to thermal stimuli, we also wondered if this behavioral stability would apply to individual animals that had recently lost neural tissue. To answer this question, we cut the organisms in half along their oral - aboral axis. Thanks to *Hydra*’s radial symmetry and regenerative abilities, these surgically altered animals repaired themselves within 48 hours, yielding a full animal with roughly half the number of neurons as the original whole (Fig. 5a-c, see Materials and Methods). By measuring neural responses to thermal stimulation both before and two days after this longitudinal sectioning, we could experimentally modulate the number of neurons in an organism and quantify resultant changes in neural activity patterns (Fig. 5b). Specifically, we generated on average a two-fold decrease in neuron count between whole and bisected *Hydra* (Fig. 5c-d); however, we found that the neural response to thermal stimulation remained stable across all stimulus temperatures from 9°C to 36°C (Fig. 5e). Namely, we found no statistically significant changes in the calcium spike rate when comparing *Hydra* before and after surgical resection (N = 5 animals per stimulus temperature). In order to account for naturally-occurring variability between animals, we performed bootstrapping on the calcium spike rates within size groups and calculated the corresponding effect size (δ) for comparison (Supplemental Fig. 3). We found that the effect sizes between the calcium spike rates of whole and bisected *Hydra* are negligible to small (δ < 0.18) for all stimulation temperatures except for 30°C, and moderate (δ = 0.41) for 30°C stimulation. These effect sizes are all small compared to the day-to-day variability within an animal (Supplemental Fig. 1) and to the change in number of neurons, which has a maximum Cliff’s delta of 1 (Fig. 5d). Thus, even when we used surgical resection to create a two-fold decrease in neuronal count, *Hydra* maintains stable encoding of thermal stimulus temperature.

**Figure 5:**
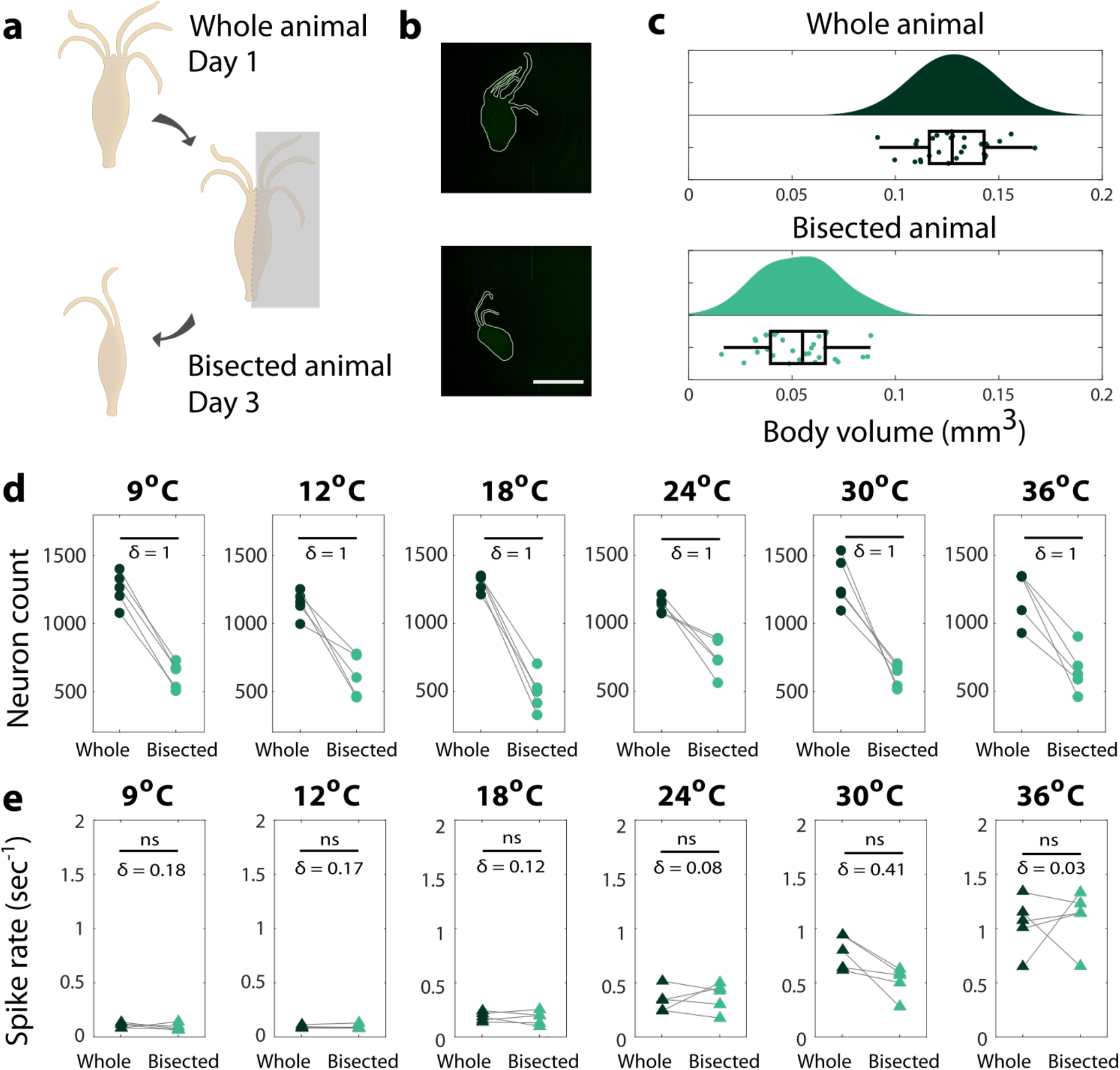
Neuron count within an animal does not affect neural response to thermal stimulation in *Hydra* vulgaris. **a** An illustration of the experiment. The whole animal is stimulated on Day 1 and the bisected animal is stimulated on Day 3 (48-52 hours after being cut). **b** Size distribution of whole and bisected *Hydra*. (N=5 for each stimulation temperature) **c** Representative images of whole *Hydra* (top panel), representative image of bisected *Hydra* (bottom panel). Scale bar = 500 μm. **d** Comparison of neuron count between whole (N=5) and bisected (N=5) *Hydra*. The dark green data points on the left correspond to the number of neurons in the whole *Hydra*, which are connected to the corresponding bisected *Hydra*, the light green points on the right. δ, the effect size using Cliff’s delta shown is calculated after bootstrapping with 100 iterations. **e** Mean spike rate comparison between whole and bisected *Hydra*. (ns = not significant, Wilcoxon signed-rank test. δ = Effect size using Cliff’s delta)

## Discussion

Unlike many invertebrate model systems like *C. elegans* and *Drosophila* that have stereotyped neural architecture with well-defined numbers of neurons, *Hydra*’s highly plastic nervous system allows us to study how behaviors like sensory-motor responses are maintained as neural circuits remodel and incorporate new neurons. The two-layer microfluidic device shown here is a well-controlled method to study this process because it allows us to simultaneously image neural activity and behavior while we deliver thermal stimuli without mechanical stimulation artifacts. Using this technology, we were able to characterize *Hydra’s* response to rapid changes in temperature. Following thermal stimulation, we found that the animal first elongates and then contracts, with the peak body length occurring approximately 20 seconds after the stimulus onset. This behavioral response to a thermal stimulus is accompanied by synchronous periodic activity in a group of neurons near the animal’s aboral end that we refer to as the TR neurons. The frequency of TR neuron activity (defined as calcium spike rates) depends on the absolute temperature of the thermal stimulus, as opposed to relative changes from the baseline culture temperature. This encoding is maintained even if the number of neurons in the animal changes by a factor of two via either natural changes to the animal’s size or by surgical resection along the oral-aboral axis.

One remaining question is the identity of the TR neurons. Their location near the animal’s foot suggests that they may be the contraction burst (CB) neurons that are associated with animal contractions [20], but more work is needed to determine if TR neurons and CB neurons are a completely overlapping neuronal subtype. This could be answered using cell type-specific promoters or other cell type labeling strategies that can be developed based on recent single-cell transcriptomic data. Among the 12 neuronal subtypes, the neurons in the peduncle have been identified to be comprised of ectodermal ganglion neurons that express *Hym-176A*, *Hym-176C* and *Hym-176D*, and ectodermal sensory and ganglion neurons expressing *RFamide preprohormone A* [22]. Considering *Hym-176C* and *Hym-176D* are only present in the peduncle, modulating neurons expressing those genes can help us get one step closer to understanding the identity of the TR neurons.

Another question is how the TR neurons maintain their thermal response properties in circuits of different sizes. It is possible that *Hydra* have homeostatic mechanisms to maintain a neural circuit architecture that has the same thermal response even as neurons are added and removed. It is also possible that the thermal response circuit is simply not affected by the overall numbers of neurons in the circuit, provided there is a critical minimum number of each constituent cell type. Alternatively, the TR response may be a cellular property that is independent of the circuit size and architecture (e.g., harnessing synaptic scaling or intrinsic homeostatic plasticity to maintain the activity set point [37]). The answer to this question could be elucidated through single cell electrophysiology, blocking cell-to-cell signaling in this circuit, or using cell-type specific promoters to alter properties of neuronal subsets. *Hydra*’s combination of extensive neural plasticity and promise for genetically targeted manipulations could help evaluate the relative contributions of cellular and ensemble-level properties to this circuit and its stability.

By studying sensory-motor transformations like this thermal response, we may also learn about fundamental properties of neural circuits that are common to many species. For example, the TR neurons show characteristics of a neural oscillator, which is a common motif across phylogeny. Such an approach could point toward conserved principles of neural circuits that appeared early in our evolutionary history.

Overall, the ability to study neural and behavioral responses to thermal stimulation in a highly regenerative animal amenable to cellular-resolution fluorescent imaging offers many advantages as a model system for uncovering how neural circuits remodel without compromising their function.

## Materials and Methods

### Microfluidic device fabrication

All microfluidic devices were fabricated using polydimethylsiloxane (PDMS) (Sylgard 184). The thermal stimulation chip is a two-layer device: bottom stimulus flow channels and a top *Hydra* immobilization chamber, separated by a glass coverslip. By separating *Hydra* from fluid flow, the animal can be rapidly heated/cooled (by flowing water at controlled temperatures through the bottom stimulus flow channels) without undesired mechanical stimulation from high flow rates (avoided due to the glass coverslip separation). To construct the double layer chip, we first fabricated the top immobilization layer by molding PDMS from a master mold adapted from a 2 mm diameter × 100 μm tall chemical perfusion chip (designed as previously described) [38]. Briefly, a circular observation chamber designed specifically for *Hydra* was patterned using soft lithography on a silicon substrate, and a ~5mm thick layer of PDMS was molded from this master mold. After punching holes for the inlet and outlet ports with a 1.5mm diameter biopsy punch, the immobilization layer was O_2_ plasma bonded to a 12mm diameter glass coverslip.

This top immobilization chip was then placed with a glass coverslip side facing down directly on the center of the master mold for the flow layer. Uncured PDMS was poured into the flow layer mold surrounding the immobilization chamber, taking care not to clog the ports in the immobilization layer. The master mold for the flow layer (adapted from Duret et al [39]) was 3D-printed with 1mm tall channels (Form 2, Flexible Resin, Formlabs). After curing the PDMS for the bottom flow layer (thereby embedding the immobilization chamber), holes were punched for inlet/outlet access for the flow layer. Finally, this double layer microfluidic chip was O_2_ plasma bonded to a 500 μm fused silica wafer (University Wafer).

After unloading *Hydra* from the device at the end of a trial (see “Loading and unloading *Hydra*”), both layers of the microfluidic device were rinsed with *Hydra* media (media adapted from the laboratory of Robert Steele), sonicated in *Hydra* media for at least 10 minutes, and soaked at room temperature in *Hydra* media. Such cleaning allowed for devices to be reused across multiple days of trials.

### Hydra strains and maintenance

All trials were conducted on transgenic lines with neuronal expression of GCaMP6s, developed with embryo microinjections by Christophe Dupre in the laboratory of Rafael Yuste. *Hydra* were cultured using the protocol adapted from the laboratory of Robert Steele (UC Irvine). *Hydra* were raised at either 18°C or 25°C in incubators, both with 12:12 hours of light:dark cycle. Animals were fed freshly hatched artemia nauplii three times a week and cleaned after approximately 4 hours with *Hydra* media. All animals used in trials were starved for a day prior to thermal stimulation experiments and were not re-used.

### Loading and unloading *Hydra*

*Hydra* were loaded into the inlet port of Hydra enclosure using a 10 mL syringe with attached tygon tubing. *Hydra* were then pulled a couple of centimeters into the tygon tubing before the tubing was inserted into the inlet port of the microfluidic device. By applying gentle pressure and gently pulsing on the plunger of the syringe, *Hydra* could be successfully loaded into the immobilization chamber without damage from mechanical shear. After the experiments, *Hydra* was removed from the device by applying pressure on the plunger of a tygon tubing-attached syringe and flushing the organism out the outlet port.

### Thermal stimulation assay

After loading *Hydra*, two programmable syringe pumps (New Era NE-500) controlled by an Arduino Mega ADK were used to drive the flow of deionized water at a rate of 6 mL/min through two inlet ports of the thermal stimulation device. The third inlet port was connected to an additional syringe (useful for removing air bubbles when initially filling the bottom stimulus flow channels prior to the start of a trial), and the device outlet port was connected to a water collection container. Fluid flow from the two pumps was heated/cooled using two in-line heaters (SC-20, Warner Instruments) regulated by a Dual Channel Bipolar Temperature Controller (CL-200A, Warner Instruments). One in-line heater supplied *Hydra*’s culture temperature during control periods (i.e. 18°C or 25°C), while the other heater supplied the desired stimulus temperature of a given trial (i.e. 9°C, 12°C, 18°C, 24°C, 25°C, 30°C, or 36°C). To compensate for heat exchange between water flowing through inlet tubing and the environment, a FLIR ONE thermal camera was used to calibrate the relationship between in-line heater temperature settings and actual temperatures of the thermal stimulation device.

Each thermal stimulation trial began with two minutes at *Hydra*’s culture temperature. Subsequently, the trial alternated between stimulus periods (single temperature for a given trial; 9°C, 12°C, 18°C, 24°C, 25°C, 30°C, or 36°C) and control periods (based on *Hydra*’s culture temperature; 18°C or 25°C). Stimulus periods always lasted for 60 seconds, while control periods varied in length between 30 and 90 seconds in 15 second increments, to help distinguish between stimulus-evoked responses and spontaneous activity. The lengths of control periods were randomly ordered and averaged to 60 seconds over the course of a trial. For a visual representation of thermal stimulation protocols, see Fig. 3a. The timing of transitions between stimulus and control periods were recorded through the Arduino’s Serial Monitor (57600 baud rate) and used to determine portions of recordings corresponding to stimulus and control periods. For experiments on size comparison of *Hydra* cultured at 18°C, 3 large and 3 small animals for each temperature (9°C, 12°C, 18°C, 24°C, 30°C, and 36°C), 36 animals in total were used over the course of 15 days. For *Hydra* cultured at 25°C, 3 animals were used at 18°C, and 4 animals at 25°C, 30°C, and 36°C over the course of 7 days.

Fluorescence imaging was conducted on a Nikon SMZ18 stereomicroscope with a SHR Plan Apo 2x objective (0.3 NA) and Andor Zyla 4.2 (16 fps, 3×3 image binning). Excitation was provided by a X-Cite Xylis XT720L at 50% intensity through a Chroma EGFP filter cube (catalog no. 49002). The start of fluid flow using Arduino and the start of recording using Andor Solis were manually synced with a maximum delay less than 1 second.

### Measuring Day-to-Day Variability in Calcium Spike Rate

To better understand the source of variability in our data we asked how much of the difference in calcium spike rates could be explained by day-to-day variability in the same animal. To answer this question, we stimulated the same whole *Hydra* twice at 36°C over a 48 hours interval (N = 3 organisms) - the same time interval between the whole and partial *Hydra* experiments. We found that while one animal maintained its spike rate with no statistically significant difference, two animals had a statistically significant decrease in spike rate, even when stimulated at the same temperature before and after the 48 hour difference (Supplemental figure 2b). Accompanying effect sizes measured using Cliff’s delta (δ) ranged from 0.04 for first animal to 0.41 and 0.61 for the latter two (overall pooled effect size of 0.35), contextualizing the relative magnitudes of observed experimental effects against typical day-to-day variability in *Hydra*.

### Longitudinal imaging of the entire nervous system

Transgenic *Hydra vulgaris* AEP expressing GFP (green fluorescent protein) in their interstitial cell lineage were used to investigate how the number of neurons varies with animal size and nutrient availability. Animals with large body size (>2 mm long) at the beginning of the study were starved for the duration of the experiment. A control group of animals with small body size (< 1mm) at the beginning of the study were fed with an excess of freshly hatched artemia nauplii three times a week. Size of the animal was defined as the length of the animal in a relaxed state (between fully contracted and fully elongated). *Hydra* media was replaced daily for all animals.

Volumetric imaging was performed using a confocal microscope (Nikon TI Eclipse) with animals immobilized in chemical perfusion microfluidic chips (see Badhiwala et al. [38]) and chemically paralyzed with 1:5000 linalool. Animals were imaged three times a week (one day after the control animals were fed) over two weeks. All images were acquired with a 10x DIC objective. The majority of the images were acquired with 1024 × 1024 pixels (x, y) resolution and 5 mm z-resolution. A lower resolution of 512 × 512 pixels and 10 mm z-resolution were used in a few cases to speed up volumetric imaging where micromotions of the animals could not be completely eliminated with chemical anesthetic (becoming increasingly important with animals starved for extended periods of time).

To quantify the total number of neurons, a maximum intensity projection image was generated from the z-stacks. After binarizing the resulting image with a user-defined threshold, individual regions (or neurons) were summed to determine the total number of neurons. However, due to anatomical overlap of multiple neurons in high-density regions such as the peduncle, future work will focus on developing transgenic animals that express fluorescent reporters in the nuclei of specific neural cell types, rather than in the cytosol of interstitial cell derived lineages (as in the animals used here).

To quantify the body size during an experiment, we binarized the maximum intensity projection image with a lower threshold than above. Filling holes in the binarized image produced the binary mask for the whole animal. Body size was calculated as the area of this binary region, and body volume was body size times the thickness of the immobilization chamber (160 μm).

The duration of the experiment was kept under two weeks, as well-fed animals can reproduce asexually by budding every 3-4 days. In fact, one of the animals in the well-fed group began forming a bud on day 10 of the experiment. Additionally, as animals are starved, they become smaller and more transparent, making them difficult to handle. In our starved group, we ‘lost’ one of the animals on day 15 as it was either too transparent or had shrunk considerably to not be discerned from the plate.

### Creating bisected *Hydra*

For the purpose of investigating whether the number of neurons affect the neural response to thermal stimulation in *Hydra*, the animals were cut along the oral - aboral axis in order to ensure we have a significantly different distribution of body size while retaining tentacles, the hypostome, body column, and peduncle. Whole, uncut *Hydra* were imaged on day 1 as the same protocol described above. Then each *Hydra* was cut along the oral-aboral axis under a dissection microscope using an X-acto knife. Any form of anaesthesia was not used in order to eliminate chemical perturbation in muscle or neural activities. The cut animals were incubated in separate 24-well compartments and were fed the day after being cut (day 2) to keep the feeding schedule consistent. After letting them recover for 48 to 50 hours, they were thermally stimulated again under the same condition they were exposed to before being cut. Five animals were used for each stimulation temperature. For the comparison of neural response between whole and bisected *Hydra*, only one of the two bisected *Hydra* was used.

### Behavioral analysis

DeepLabCut [34] along with custom MATLAB code was used to quantify *Hydra*’s behavioral responses to thermal stimuli. 20 frames per video were extracted for manual tracking according to k-means clustering on DeepLabCut. *Hydra* in each frame was then manually annotated with the locations of the basal disc (aboral end), left side, center, and right side of the body, and hypostome (oral end). Two corners of the immobilization chamber were additionally annotated, providing a known distance (2 mm) to convert pixel measurements to micrometer measurements. The annotated dataset was used as training data for DeepLabCut. After evaluating the model, the videos were analyzed using the trained model which yielded the coordinates of the seven points listed above for every frame (total of 14,600 frames) for every video, along with annotated videos. With the coordinates obtained from DeepLabCut annotations, body width was calculated as the sum of the distance from the left side to the middle of the body and the distance from the middle to the right side of the body, and body length was calculated as the sum of the distance from the hypostome to the middle of the body and the distance from the middle of the body to the basal disc (Fig. 2b, bottom panel) with a custom MATLAB code.

### Analysis of neural activity in the peduncle of *Hydra*

To determine the timing and frequency of calcium spikes in *Hydra*’s peduncle oscillator, fluorescence microscopy recordings from thermal stimulation trials of neuronal GCaMP6s *Hydra* (see “Thermal Stimulation Assay”) were processed in Fiji (ImageJ) and in MATLAB. For each frame of the recording, Fiji was used to calculate the average intensity of a ROI encompassing *Hydra*’s peduncle and aboral regions (same ROI for the entire recording). In MATLAB, this intensity trace was smoothed (5 data point span), and the built-in findpeaks function was used to determine spike locations based on their prominence in the intensity traces. Spurious spikes (e.g. from measurement noise) were eliminated by additionally testing that each identified peak was the only local maximum exceeding a threshold prominence value within a narrow window of the peak.

Raster plots were then directly produced from the times of peduncle oscillator spikes during each stimulus period and the surrounding 30 seconds at culture temperature. Peristimulus time histograms defined as calcium firing rate were calculated by counting the number of spikes across all stimulus periods at a given stimulation temperature in a 10-second sliding window, then normalizing by the number of stimulus periods and the length of the window. Instantaneous calcium spike rates for a given thermal stimulus temperature were defined and calculated as the inverse of interspike intervals during stimulus periods.

## Acknowledgements

We would like to thank Dr. Christophe Dupre and Dr. Rafael Yuste (Columbia University), Dr. Celina Juliano (UC Davis) and Dr. Rob Steele (UC Irvine) for sharing transgenic *Hydra* lines, and Dr. Juliano for comments on the manuscript. This work was supported in part by the NSF EDGE.

C.N.T. is funded by a Fannie and John Hertz Foundation Fellowship and by a National Science Foundation Graduate Research Fellowship. K.N.B. is funded by training fellowships from the Keck Center of the Gulf Coast Consortia on the NSF Integrative Graduate Education and Research Traineeship (IGERT): Neuroengineering from Cells to Systems 1250104.

## Author Contributions

C.N.T. and S.K. performed and analyzed *Hydra* thermal stimulation experiments. K.N.B. performed and analyzed longitudinal imaging experiments on *Hydra*’s number of neurons as a function of body size. C.N.T., K.N.B., and B.W.A. designed, prototyped, and fabricated the microfluidic device. C.N.T., S.K., and B.W.A. implemented microcontroller automation of thermal stimulation experiments. C.N.T., J.T.R. supervised the research. C.N.T., S.K., K.N.B., and J.T.R. co-wrote the paper. All authors read and commented on the manuscript.

## Supplemental Information

**Supplemental Figure 1:**
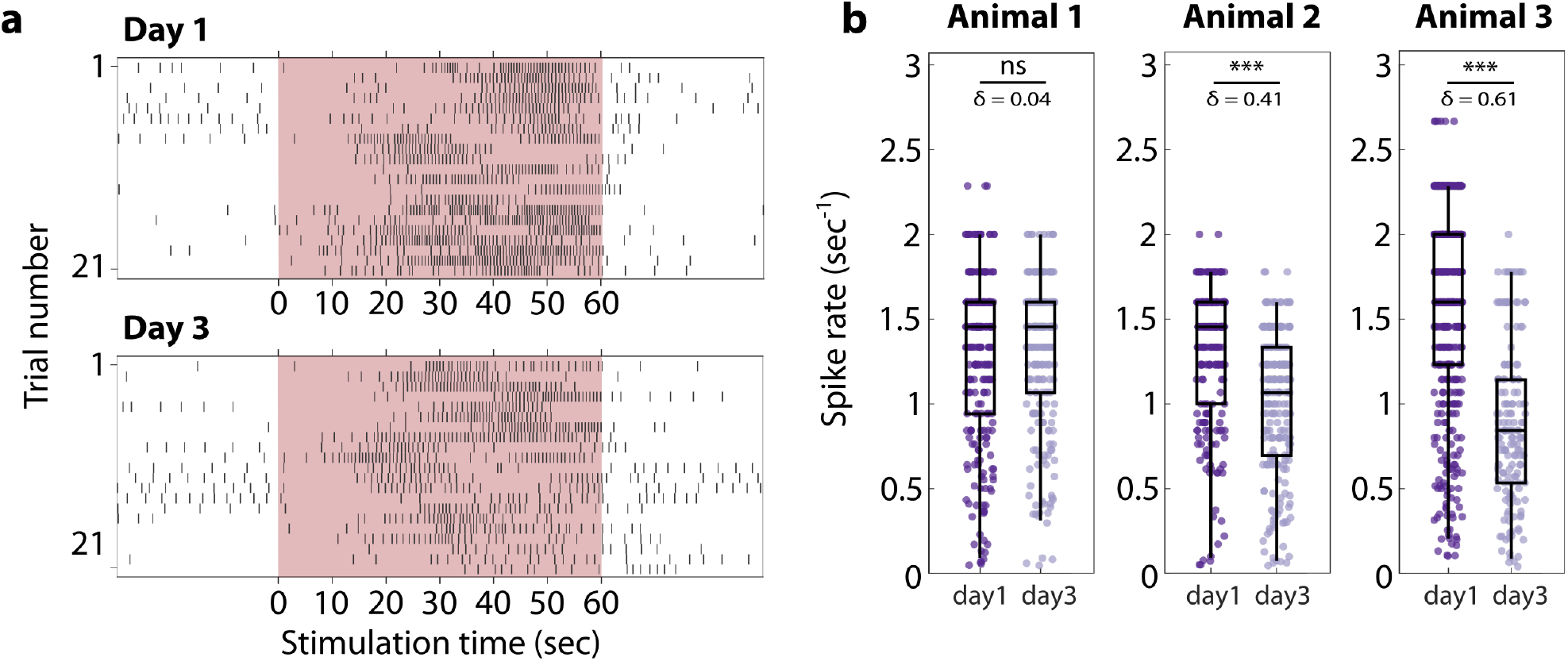
Day-to-day variance in intra-animal neural activity. **a** Raster plot of *Hydra* stimulated at 36°C on Day 1 and Day 3. **b** Calcium spike rate comparison between day1 and day3 for individual animals (Mann-Whitney U test, ns = not significant, *** p<0.0001, δ = Effect size using Cliff’s delta)

**Supplemental Figure 2:**
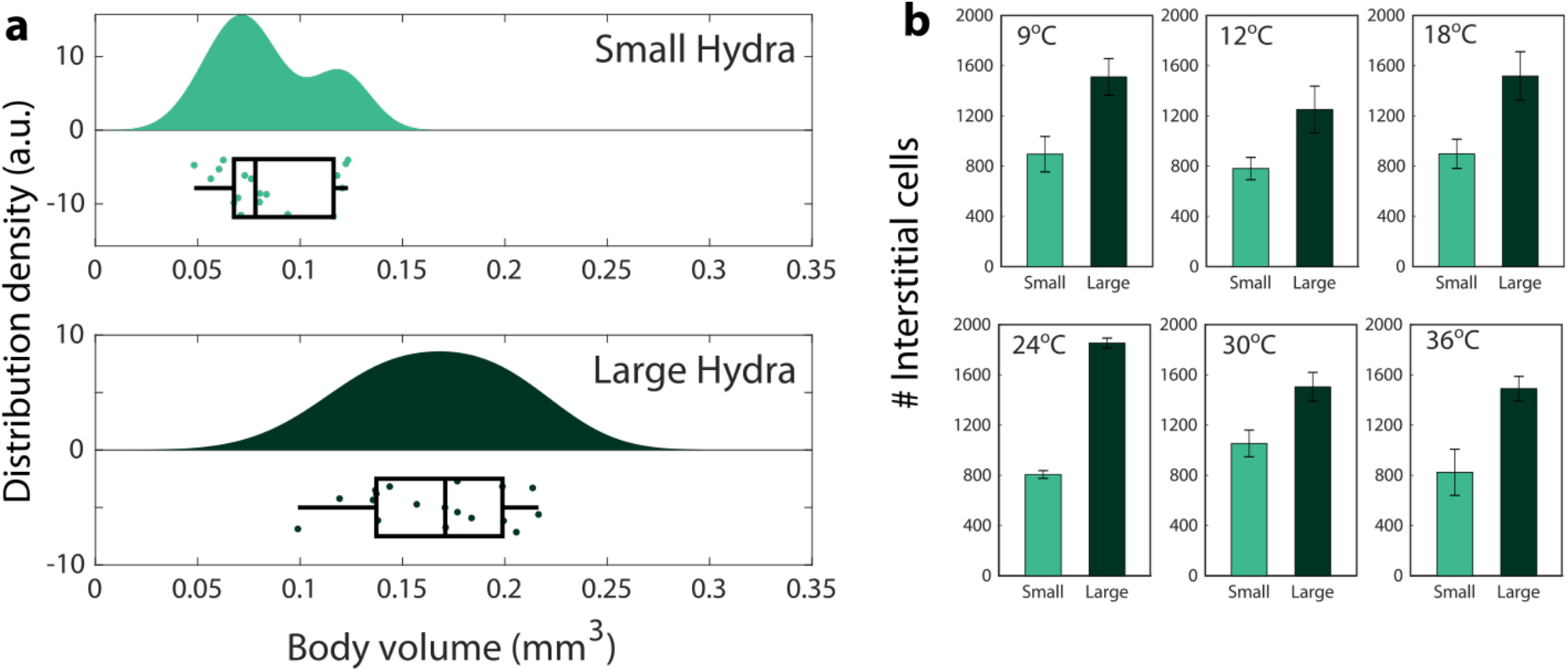
Differences in neuron counts due to *Hydra*’s natural size variations. **a** Size distribution of small and large *Hydra*. **b** Comparison of neuron count of small and large *Hydra* groups at each stimulation temperature. (N=3)

**Supplemental Figure 3:**
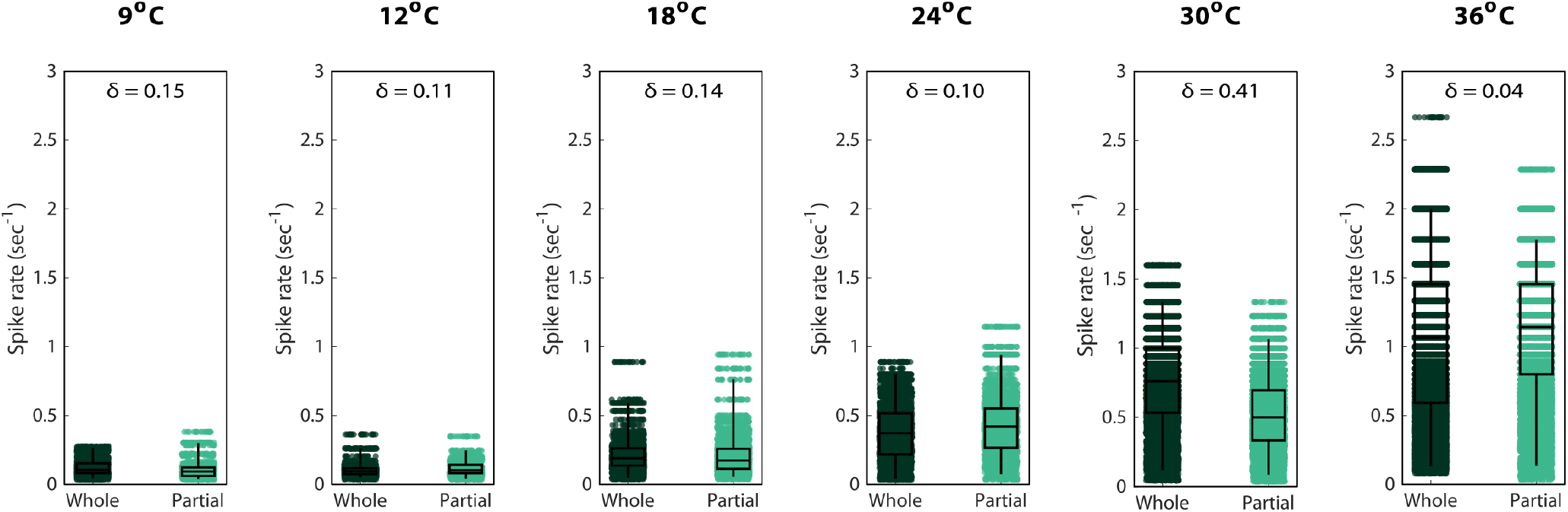
Bootstrapping analysis on spike rates of whole and bisected Hydra. Each animal was stimulated 7 times in one experiment (see Materials and Methods), with 5 animals per stimulation temperature. For bootstrapping, we pooled calcium spike rates from 7 stimulation periods across the aforementioned 5 animals - at least one period from each of the animals - to create one artificial animal. Bootstrapping was done 100 times to create the calcium spike rates shown in this figure

**Supplemental Video 1:** *Hydra* that was cultured at 18°C undergoing a thermal stimulation assay, with non-stimulus periods at 18°C and stimulus periods at 30°C. The yellow trace indicates fluorescence changes in an ROI encompassing TR neurons, and orange-brown rectangles indicate stimulus periods.

**Supplemental Video 2:** *Hydra* that was cultured at 18°C undergoing a thermal stimulation assay, with non-stimulus periods at 18°C and stimulus periods at 18°C. The yellow trace indicates fluorescence changes in an ROI encompassing TR neurons, and orange-brown rectangles indicate stimulus periods.

